# 5,6-dimethylxanthenone-4-acetic acid (DMXAA), a Partial STING Agonist, Competes for Human STING Activation

**DOI:** 10.1101/2023.12.07.570548

**Authors:** Burcu Temizoz, Takayuki Shibahara, Kou Hioki, Tomoya Hayashi, Kouji Kobiyama, Michelle Sue Jann Lee, Naz Surucu, Erdal Sag, Atsushi Kumanogoh, Masahiro Yamamoto, Mayda Gursel, Seza Ozen, Etsushi Kuroda, Cevayir Coban, Ken J. Ishii

**Affiliations:** Division of Vaccine Science, Department of Microbiology and Immunology, The Institute of Medical Science, The University of Tokyo, Tokyo, Japan; International Vaccine Design Center (VDesC), The Institute of Medical Science (IMSUT), The University of Tokyo, Tokyo, Japan; Department of Respiratory Medicine and Clinical Immunology, Graduate School of Medicine, Osaka University, Osaka, Japan; Division of Malaria Immunology, Department of Microbiology and Immunology, The Institute of Medical Science (IMSUT), The University of Tokyo, Tokyo, Japan; Department of Biological Sciences, Middle East Technical University (METU), Ankara, Turkey; Department of Pediatric Rheumatology, Hacettepe University, Ankara, Turkey; Department of Immunoparasitology, Division of Infectious Disease, Research Institute for Microbial Diseases, Osaka University, Osaka, Japan; Izmir Biomedicine and Genome Center, Izmir, Turkey; Department of Immunology, School of Medicine, Hyogo Medical University, Hyogo, Japan; Immunology Frontier Research Center (IFReC), Osaka University, Osaka, Japan; Core Research for Evolutional Science and Technology (CREST), Japan Science and Technology Agency (JST), Tokyo, Japan

**Keywords:** STING, partial agonist, DMXAA derivative, SAVI, HHMX

## Abstract

5,6-dimethylxanthenone-4-acetic acid (DMXAA) is a mouse-selective stimulator of interferon gene (STING) agonist exerting STING-dependent anti-tumor activity. Although DMXAA can not fully activate human STING, DMXAA reached phase III in lung cancer clinical trials. How DMXAA is effective against human lung cancer is completely unknown. Here, we show that DMXAA is a partial STING agonist interfering with agonistic STING activation, which may explain its partial anti-tumor effect observed in humans, as STING was reported to be pro-tumorigenic for lung cancer cells with low antigenicity. Furthermore, we developed a DMXAA derivative—3-hydroxy-5-(4-hydroxybenzyl)-4-methyl-9*H*-xhanthen-9one (HHMX)—that can potently antagonize STING-mediated immune responses both in humans and mice. Notably, HHMX suppressed aberrant responses induced by *STING* gain-of-function mutations causing STING-associated vasculopathy with onset in infancy (SAVI) in *in vitro* experiments. Furthermore, HHMX treatment suppressed aberrant STING pathway activity in peripheral blood mononuclear cells from SAVI patients. Lastly, HHMX showed a potent therapeutic effect in SAVI mouse model by mitigating disease progression. Thus, HHMX offers therapeutic potential for STING-associated autoinflammatory diseases.

## Introduction

Stimulator of interferon genes (STING) is one of the key molecules at the intersection of various cytosolic nucleic acid-sensing pathways, including cyclic GMP-AMP synthase (cGAS), DEAD-box helicase family 41, and interferon gamma inducible protein 16 (1, 2). One of the most commonly studied cytosolic sensor, namely, cGAS, can detect cytosolic DNA or RNA:DNA hybrids to catalyze the production of the noncanonical cyclic dinucleotide 2′3′cGAMP (or mammalian cGAMP), which can directly bind to and activate the STING-TBK1-IRF3 pathway, resulting in type I Interferon (IFN) and proinflammatory cytokine production (1–3). In addition to 2′3′cGAMP, small molecule 5,6-dimethylxanthenone-4-acetic acid (DMXAA) or cyclic dinucleotides (CDNs) of prokaryotic origin, such as c-di-AMP, c-di-GMP, and 3′3′cGAMP (canonical), are also capable of activating STING via direct binding (4–6).

Several reports have demonstrated that STING agonists function as potent vaccine adjuvants by eliciting antigen-specific T and B cell responses (4,7,8). Along with the adjuvant effects of STING agonists, potent anti-tumor immunity can also be elicited by STING agonists via mechanisms involving the induction of type I IFN and CTL responses (9–11). By contrast, the inappropriate detection of the cytosolic host DNA leaking from damaged cell nuclei or the continuous activation of the STING pathway leads to the development of autoinflammatory conditions such as systemic lupus erythematosus (SLE) (11). More recently, gain-of-function mutations in STING were identified as the cause of a severe autoinflammatory condition known as STING-associated vasculopathy with onset in infancy (SAVI), distinguished by interferonopathy and subsequent systemic inflammation, skin lesions, and interstitial lung disease (12). Moreover, mutations in the DNase enzyme three prime exonuclease 1 (TREX1) were associated with the development of autoinflammatory conditions such as Aicardi–Goutieres syndrome (AGS) and SLE, culminating in autoinflammation-mediated lethality, which is dependent on aberrant STING pathway activation as a result of the inability of the cells to clear DNA from dying cells (13,14). Together, these data emphasize the importance of the STING pathway for the development of autoinflammatory diseases and highlight the necessity of STING antagonist-based immunotherapies.

The chemotherapeutic agent DMXAA exerts potent anti-tumor activity in mouse tumor models through various mechanisms, including the disruption of tumor vasculature, nitric oxide induction from tumor-associated macrophages, and the induction of tumoricidal cytokines such as TNFα and type I IFNs, the latter in a STING-dependent manner (13, 15 and 16). However, the anti-tumor activity of DMXAA was found to be limited in humans after it failed in phase III clinical trials of non-small cell lung cancer (NSCLC) (15). Subsequently, this failure was attributed to the fact that DMXAA could activate only the mouse, but not the human STING pathway (16). However, it remains unclear how DMXAA was able to enter phase III trials, if it cannot activate the human STING pathway. By contrast, Ahn et al. demonstrated that chronic inflammation triggers tumorigenesis via STING (17). As inflammation is one of the key factors for inducing tumorigenesis, in addition to the fact that STING was shown to promote the tumorigenesis of lung cancer tumors with low antigenicity via indoleamine 2,3 dioxygenase (IDO) activation, it would be reasonable to hypothesize that the partial anti-tumor effect of DMXAA is due to its suppressive activity on STING-induced inflammatory responses in humans (18,19).

Based on these evidence, we propose that DMXAA may be an antagonist or partial agonist for the human STING pathway, suppressing STING-induced inflammatory responses in humans. Thus, we investigated the suppressive effect of DMXAA using human peripheral blood mononuclear cells (hPBMCs) and THP1 dual reporter cells in vitro and found that DMXAA acted as a partial human STING agonist competing with agonistic STING activation. Moreover, we also developed a novel DMXAA derivative—3-hydroxy-5-(4-hydroxybenzyl)-4-methyl-9H-xhanthen-9one (HHMX) —and tested the therapeutic potential of HHMX not only in J774 dual reporter cells expressing STING constructs with various SAVI mutations but also in PBMCs isolated from a SAVI patient with heterozygous STING N154S mutation (human counterpart of mouse N153S mutation). Finally, in vivo studies using SAVI mice were conducted by using HHMX to test its therapeutic potential. Overall, our results revealed key mechanisms of action of DMXAA in humans and revealed a novel DMXAA derivative, HHMX, which can be a potential therapeutic agent for STING-mediated inflammatory diseases.

## Results

### DMXAA is a partial agonist for human STING, interfering with agonistic STING activation

Inflammation is a well-known factor involved in tumorigenesis (20). Moreover, considering the findings of Ahn et al. regarding the STING-dependency of inflammation-driven tumorigenesis (17), we hypothesized that the suppressive activity of DMXAA on the STING pathway in human cells could be the reason for it demonstrating partial anti-tumor activity in humans and managing to reach phase III in clinical trials of NSCLC. Therefore, we hypothesized that DMXAA acts as an antagonist or partial agonist for human STING to suppress STING-induced immune responses. To verify this hypothesis, we stimulated hPBMCs with several STING agonists after DMXAA pretreatment and screened several cytokines induced by STING pathway activation. Interestingly, we found that DMXAA exerted a suppressive effect on STING-induced type I and II IFN production, whereas STING-induced IL-6 production remained unaffected (Figure 1A). Furthermore, this suppressive effect of DMXAA was not due to cytotoxicity as DMXAA did not cause considerable cell death compared to the control treatment (Figure S1A). It is worth noting that variable cytokine responses to different STING agonists by each PBMC donor is most likely due to STING polymorphisms, as reported previously (21). In addition, the multiplex cytokine analysis of hPBMC supernatants from additional two healthy donors by 25-plex ProcartaPlex kit indicated that, DMXAA had a partial suppressive effect on STING agonist-induced cytokine and chemokine responses, including IP-10, granzyme B, MIP-1β, and MCP1 (Figure S1B), but not remarkably on others including eotaxin, CXCL13, IL-1a, IL-1b, IL1-RA, IL-2, IL-4, IL-5, IL-6, IL-7, IL-8, IL-9, IL-10, IL-12p40, IL-13, IL-15, IL-17A, MIP-1a, MIG, MCP-1, TNFa, sCD40L and sFasL (Figure S1C).

**Figure 1:**
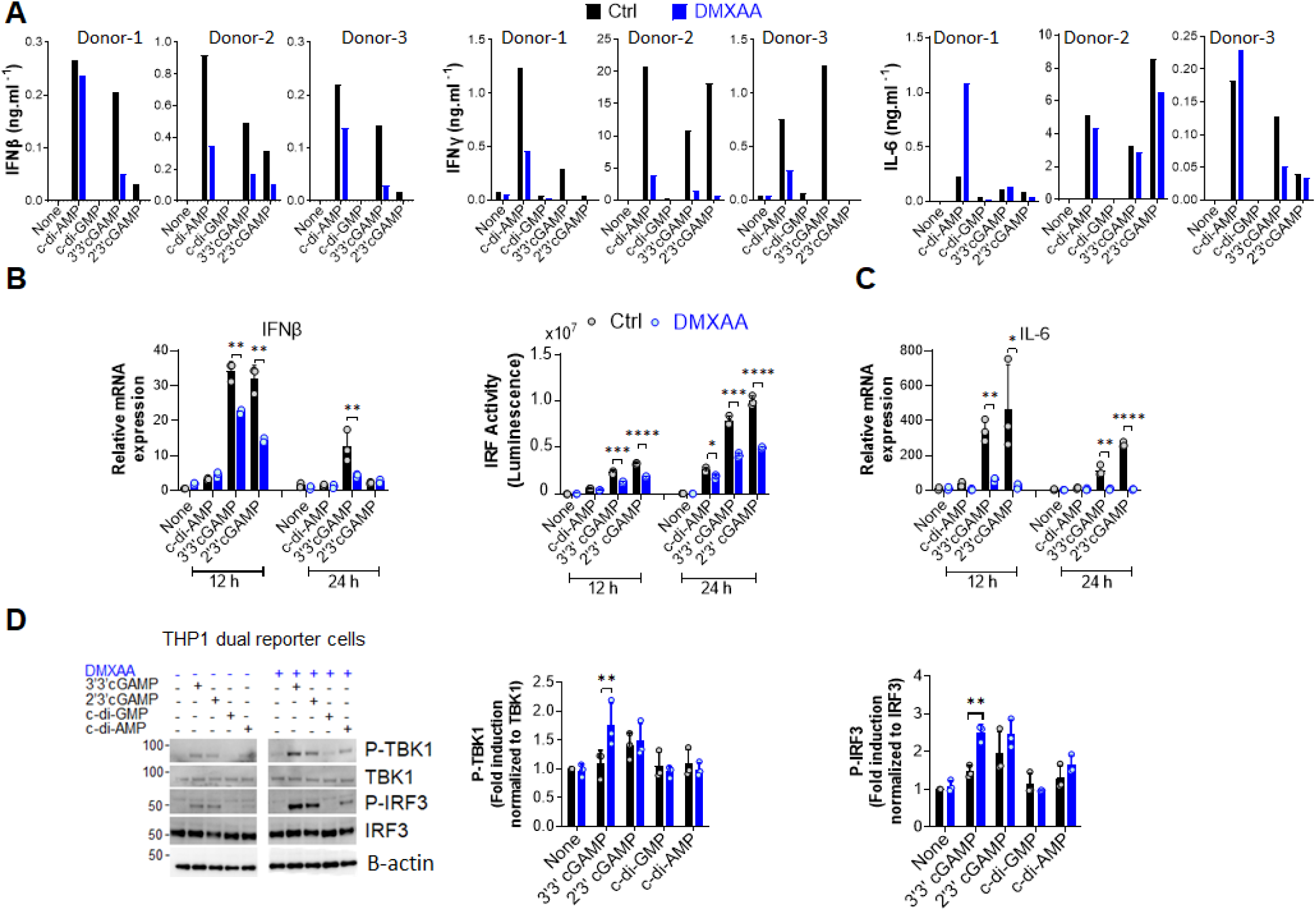
DMXAA is a partial agonist for human STING interfering with agonist-mediated STING activation. **(A)** Fresh human PBMCs from three healthy donors (n=3) were stimulated with the indicated CDNs (10 µg/ml) for 24 h after 90 minutes of DMXAA (100 µg/ml) pretreatment. IFNβ, IFNγ, and IL-6 production was measured using ELISA. Data from each individual from two independent experiments are shown as bar graphs. **(B and C)** PMA-differentiated THP1 dual reporter cells were pretreated with DMXAA (100 µg/ml) for 90 min and stimulated with the indicated CDNs (10 µg/ml) for 12 or 24 h in three independent experiments. After RNA isolation and cDNA synthesis, the mRNA expression levels of IFNβ **(B)** and IL-6 **(C)** were measured using RT-PCR. 18S ribosomal RNA was used as an internal control. IRF activity **(B)** was determined by measuring secreted luciferase activity in the supernatants. Representative data from three independent experiments are shown as the mean ± SD of triplicates (*p < 0.05, **p < 0.01, ***p < 0.001, ****p < 0.0001, Student’s t test). **(D)** PMA-differentiated THP1 dual reporter cells were pretreated with DMXAA (100 µg/ml) for 90 min and stimulated with the indicated CDNs (10 µg/ml) for 3 h in three independent experiments. P-TBK1, TBK1, P-IRF3, and IRF3, levels in the cell lysates were detected using western blotting and the blots from a representative experiment were shown. Β-actin was used as loading control. Fold induction levels for P-TBK1 and P-IRF3 relative to total TBK1 and IRF3 from three independent experiments were plotted as mean ± SD (n=3) (*p < 0.05, **p < 0.01, Student’s t test).

Next, to examine the mechanisms underlying this suppressive effect in human cells, we stimulated phorbol 12-myristate 13-acetate (PMA)-differentiated THP1 dual reporter cells with several CDNs after DMXAA pretreatment and measured STING-induced immune responses, namely the activation of the TBK1-IRF3-type I IFN and TBK1-NF-κB-IL-6 axes. In agreement with hPBMC data, DMXAA significantly suppressed STING-induced IFNβ production at the mRNA expression level and IRF promoter activation at both 12 and 24 h time points (Figure 1B). In addition, a significant suppressive activity of DMXAA on STING-induced IL-6 production was observed at the mRNA expression level at both time points (Figure 1C). However, no suppressive effect of DMXAA was observed at the TBK1, or IRF3 phosphorylation levels (Figure 1D). Indeed, DMXAA significantly enhanced 3’3’cGAMP-induced IRF3 and TBK1 phosphorylations (Figure 1D), suggesting that DMXAA may be a partial agonist for human STING competing with agonist-mediated STING activation.

### Kinetics of the suppressive effects of DMXAA in THP1 dual reporter cells

To further investigate the mechanisms of DMXAA-mediated STING pathway antagonism, we performed a dose–response experiment in PMA-differentiated THP1 dual reporter cells. We found that the significant suppression by DMXAA for IRF activity was achieved at 100 µg/ml dose while significant suppressive effect of DMXAA on STING-induced type I IFN production could be observed at a lower concentrations less than 30 µg/ml dose (Figure 2B), suggesting that relatively higher doses of DMXAA can antagonize STING-mediated immune responses.

**Figure 2:**
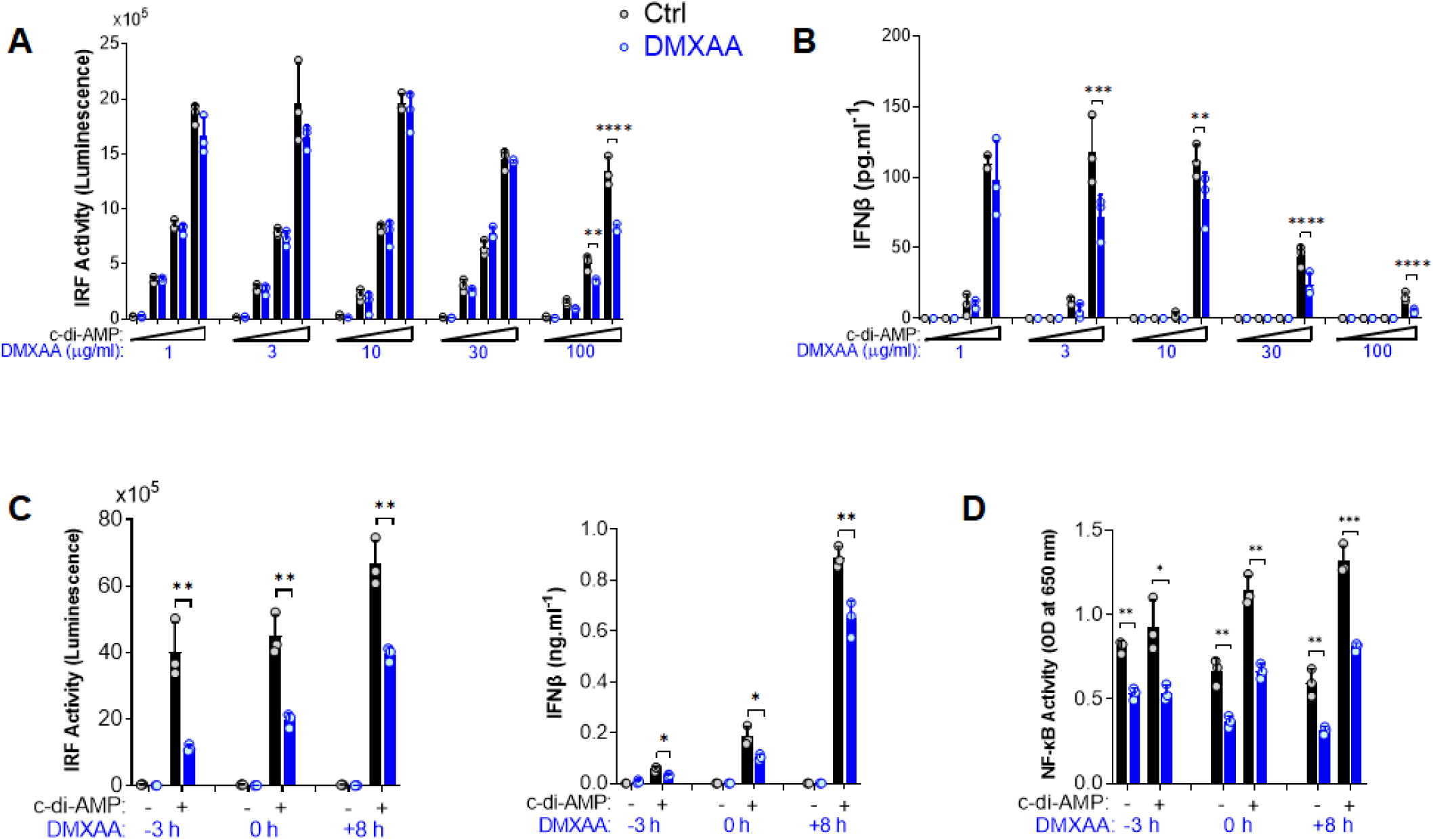
Kinetics of the suppressive effect of DMXAA in THP1 dual reporter cells. **(A)** PMA-differentiated THP1 dual reporter cells were stimulated with various concentrations of DMXAA (1, 3, 10, 30, or 100 µg/ml) together with increasing concentrations of c-di-AMP (3, 10, or 30 µg/ml) for 24 h in three independent experiments. IRF activity was determined by measuring secreted luciferase activity in the supernatants. Representative data from three independent experiments are shown as the mean ± SD of triplicates (**p < 0.01, ****p < 0.0001, One-way ANOVA with Sidak’s multiple comparison test). **(B)** IFNβ production was measured using ELISA. Representative data from three independent experiments are shown as the mean ± SD of triplicates (**p < 0.01, ***p < 0.001****, p < 0.0001, One-way ANOVA with Sidak’s multiple comparison test). **(C)** PMA-differentiated THP1 dual reporter cells were stimulated with DMXAA (100 µg/ml) at the indicated time points (‒3, 0, or 8 h) together with c-di-AMP (30 µg/ml) for 24 h in three independent experiments. IRF activity was determined by measuring secreted luciferase activity in the supernatants. IFNβ production was measured using ELISA. Representative data from three independent experiments are shown as the mean ± SD of triplicates (*p < 0.05, **p < 0.01, Student’s t test). **(D)** NF-κB activity was determined by measuring secreted embryonic alkaline phosphatase (SEAP) activity in the supernatant. Representative data from three independent experiments are shown as the mean ± SD of triplicates (*p < 0.05, **p < 0.01, ***p < 0.01, Student’s t test).

After determining the optimum dose for the antagonistic activity of DMXAA as 100 µg/ml, we next performed a time course experiment by varying the time of DMXAA addition to the culture. As a result, we found that the optimal dose of DMXAA could significantly suppress STING-induced IRF and type I IFNs, and NF-κB even after 8 h of STING pathway activation (Figure 2C and D). These findings suggest that either DMXAA is a partial STING agonist strongly competing for agonistic STING binding, or DMXAA antagonizes STING activation by regulating the STING pathway at the post-transcriptional levels downstream of TBK1, IRF3, and NF-κB.

### DMXAA derivative HHMX demonstrates potent suppression of STING-induced immune responses in both humans and mice

Since DMXAA had a slight and inconsistent inhibition on STING-mediated immune responses in hPBMCs (Figure 1A and S1B), we opted to develop DMXAA derivatives using the chemical structure of DMXAA to generate robust STING antagonists. After screening 69 chemical DMXAA derivatives, we identified several suppressor reagents. Among them, only HHMX (or #64) showed potent suppressive effects on STING-induced type I IFN responses without causing significant cell death in hPBMCs (Figure S2A and B). Consequently, we decided to focus solely on the DMXAA derivative HHMX for further experiments (Figure 3A).

**Figure 3:**
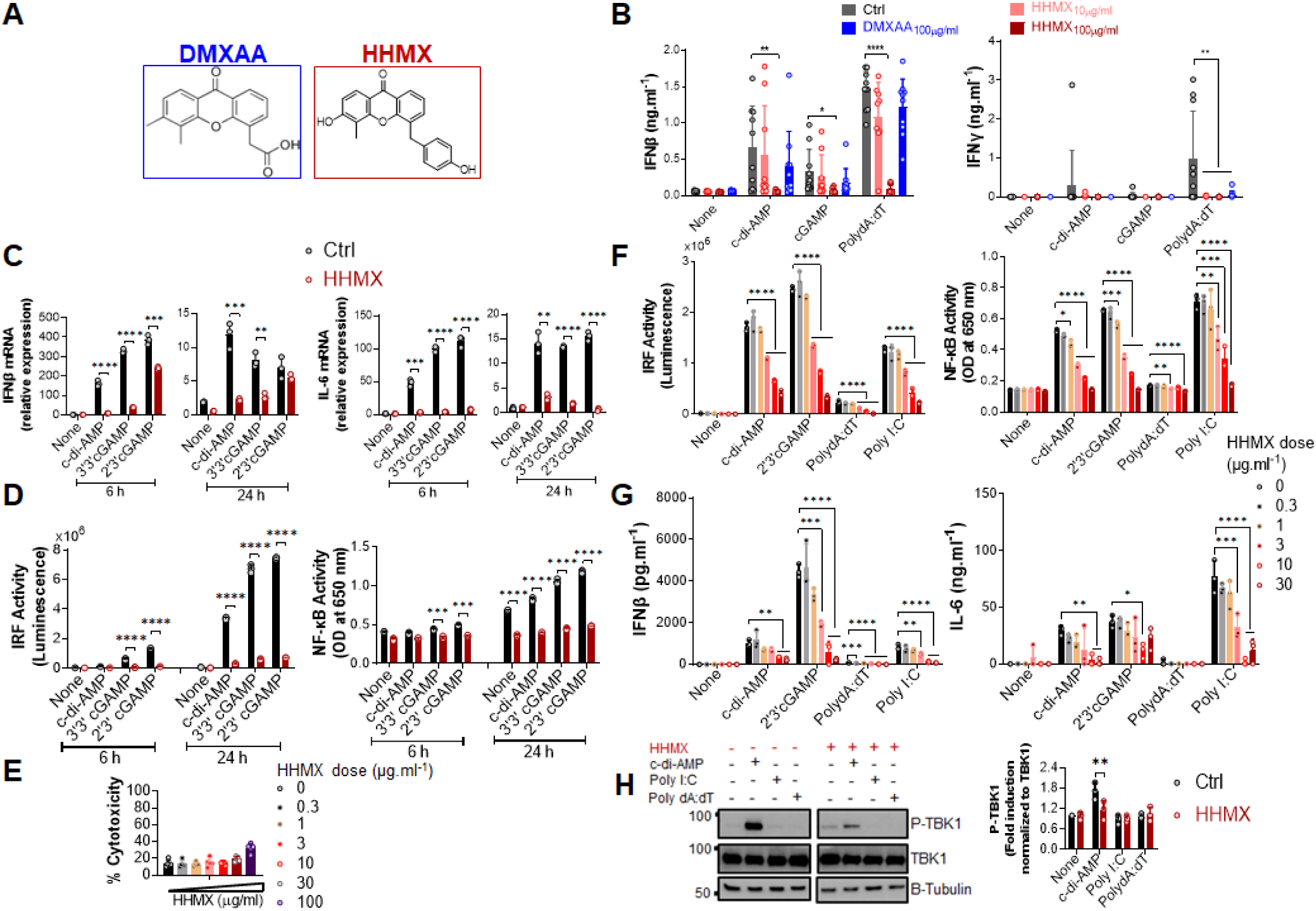
DMXAA derivative HHMX is a potent suppressor of STING-induced immune responses not only in human but also in mice. **(A)** Chemical structures of DMXAA and its derivative HHMX. **(B)** Fresh human PBMCs from healthy donors (n=10) were stimulated with the indicated CDNs (10 µg/ml) or transfected with PolydA:dT (1 µg/ml) for 24 h after 90 min of DMXAA (100 µg/ml) pretreatment. IFNγ and IFNβ productions were measured using ELISA (*p < 0.05, **p < 0.01, ****p < 0.0001, Student’s t test). **(C)** PMA-differentiated THP1 dual reporter cells were stimulated with HHMX (30 µg/ml) together with the indicated CDNs (10 µg/ml) for 6 or 24 h in three independent experiments. After the RNA isolation and cDNA synthesis, the mRNA expression levels of IFNβ and IL-6 were measured using RT-PCR. 18S ribosomal RNA was used as an internal control. Representative data from three independent experiments are shown as the mean ± SD of triplicates (**p < 0.01, ***p < 0.001, ****p < 0.0001, Student’s t test). **(D)** IRF activity was determined by measuring secreted luciferase activity in the supernatants, whereas NF-κB activity was determined by measuring secreted embryonic alkaline phosphatase (SEAP) activity in the supernatant. Representative data from three independent experiments are shown as the mean ± SD of triplicates (***p < 0.001, ****p < 0.0001, Student’s t test). **(E)** Cell death was measured using lactate dehydrogenase (LDH) release assay (from five independent experiments, n=5, biological replicates) according to manufacturer’s instructions. **(F)** J774 dual reporter cells were stimulated with increasing concentrations of HHMX together with the indicated CDNs (10 µg/ml), PolydA:dT, or Poly I:C (1 µg/ml) for 24 h in three independent experiments. IRF activity was determined by measuring secreted luciferase activity in the supernatants. NF-κB activity was determined by measuring secreted embryonic alkaline phosphatase (SEAP) activity in the supernatant. Data are shown as the mean ± SD from three independent experiments (n=3) (*p < 0.05, **p < 0.01, ***p < 0.001, ****p < 0.0001, One-way ANOVA with Sidak’s multiple comparison test). **(G)** IFNβ and IL-6 concentrations in the supernatants were determined using ELISA. Data are shown as the mean ± SD from three independent experiments (n=3) (*p < 0.05, **p < 0.01, ***p < 0.001, ****p < 0.0001, One-way ANOVA with Sidak’s multiple comparison test). **(H)** J774 dual reporter cells were stimulated with HHMX (30 µg/ml) together with the indicated c-di-AMP (10 µg/ml), PolydA:dT, or Poly I:C (1 µg/ml) for 3 h in three independent experiments. P-TBK1, TBK1 and β-tubulin in the cell lysates were detected using western blotting and the blots from a representative experiment were shown. Β-tubulin was used as loading control. Fold induction levels for P-TBK1 relative to total TBK1 from three independent experiments were plotted as mean± SD. Data are representative of three independent experiments (n=3) (**p < 0.01, One-way ANOVA with Sidak’s multiple comparison test).

To directly compare the suppressive effects of HHMX and DMXAA, PBMCs from 10 different donors were stimulated with the indicated CDNs or transfected with PolydA:dT with or without treatment of DMXAA or HHMX. HHMX was found to exhibit more potent suppressive activity on STING-induced type I and II IFN responses in comparison to DMXAA, as it almost completely shuts down cytokine production in all PBMCs tested (Figure 3B) without affecting cell viability (Figure S2B).

Next, to investigate the mechanisms of HHMX-mediated suppression in human cells, we stimulated PMA-differentiated THP1 dual reporter cells with several CDNs along with HHMX treatment and measured the STING-induced immune responses. In correlation with the hPBMC data, HHMX significantly and profoundly suppressed STING-induced IRF and NF-κB promoter activation, type I IFN and IL-6 responses at the mRNA level (Figure 3 C and D), and type I IFN production at the protein level (Figure S2C), without affecting cell viability (Figure 3E and S2B and D).

Because of the strong suppressive effect of HHMX on the human STING pathway, we next examined the suppressive effect of HHMX in mouse cells using J774 dual reporter cells. Interestingly, we found that HHMX showed a significant dose-dependent suppressive effect on STING-(activated by CDNs and poly dA:dT) or TBK1-mediated (activated by Poly I:C downstream of TLR3 and/or retinoic acid-inducible protein I (RIG-I) and melanoma differentiation-associate gene 5 (MDA5) (22)) IRF and type I IFN responses, as well as NF-κB and IL-6 responses in mouse cells (Figure 3F and G). Moreover, HHMX was found to strongly inhibit STING-induced TBK1 phosphorylation (Figure 3H). Taken together, these data indicated that HHMX could act as a potent inhibitor for STING and TBK1-related pathways not only in humans but also in mice.

### Kinetics of the suppressive effect of HHMX in THP1 dual reporter cells

The suppressive effect of HHMX was also analyzed by performing dose–response and time course experiments in THP1 dual reporter cells. HHMX showed a dose-dependent but more potent suppressive effect on STING-induced IRF and type I IFN responses than DMXAA, with even low doses of HHMX (e.g. 1 and/or 3 µg/ml), demonstrating a significant suppressive effect, in contrast to DMXAA, which showed the most robust and significant suppressive effect on STING-induced IRF or NF-κB responses at a dose of 100 µg/ml (Figure 4A). Furthermore, time course experiments using 30 µg/ml of HHMX in PMA-differentiated THP1 dual reporter cells showed that HHMX could potently suppress STING-induced IRF, type I IFN, and NF-κB responses even after 3 h of STING agonist addition (Figure 4B and C). Thus, we have come to the conclusion that the DMXAA derivative HHMX is a potent antagonist of the STING pathway in both the human and the mouse.

**Figure 4:**
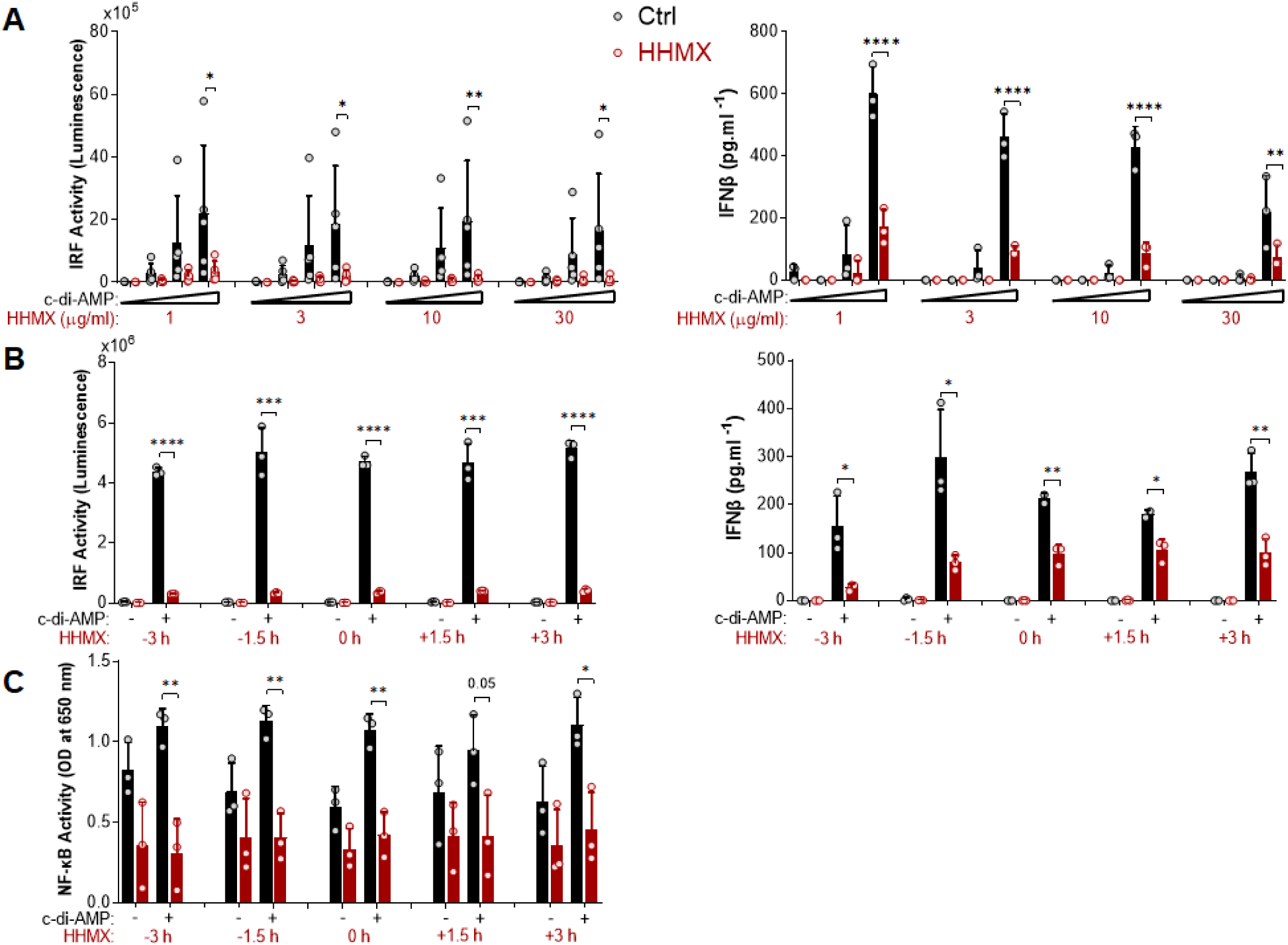
Kinetics of the suppressive effect of HMMX in THP1 dual reporter cells. **(A)** PMA-differentiated THP1 dual reporter cells were stimulated with various concentrations of HHMX (1, 3, 10, or 30 µg/ml) together with increasing concentrations of c-di-AMP (3, 10, or 30 µg/ml) for 24 h in five independent experiments. IRF activity was determined by measuring secreted luciferase activity in the supernatants Data are shown as the mean ± SD of five independent experiments (n=5). IFNβ concentration in the supernatants was determined using ELISA. Data are shown as the mean ± SD from three independent experiments (n=3) (*p < 0.05, **p < 0.01, ****p < 0.0001, One-way ANOVA with Sidak’s multiple comparison test). **(B)** PMA-differentiated THP1 dual reporter cells were stimulated with HHMX (30 µg/ml) at indicated the time points (‒3, ‒1.5, 0, 1.5, or 3 h) together with c-di-AMP (30 µg/ml) for 24 h in three independent experiments. IRF activity was determined by measuring secreted luciferase activity in the supernatants. Representative data from three independent experiments are shown as the mean ± SD of triplicates from one experiment (n=3, technical replicates). IFNβ concentration in the supernatants was determined using ELISA. Representative data from three independent experiments are shown as the mean ± SD of triplicates from one experiment (*p < 0.05, **p < 0.01, ***p < 0.001, ****p < 0.0001, Student’s t test.) **(C)** NF-κB activity was determined by measuring secreted embryonic alkaline phosphatase (SEAP) activity in the supernatant. Data are shown as the mean ± SD from three independent experiments (n=3) (*p < 0.05, **p < 0.01, Student’s t test.)

### HHMX shows therapeutic potential in STING-mediated autoinflammatory diseases

Since our data showed that the DMXAA derivative HHMX is a potent antagonist of the STING pathway, we decided to choose to test the therapeutic potential of HHMX for SAVI. To this end, we transiently transfected J774 dual reporter cells with constructs expressing WT STING or STING with various SAVI-associated mutations, including V146L, N153S, V154M, C205Y, R280Q, and R2833G (12,13, 29), and measured STING-mediated immune responses with or without HHMX treatment. As expected, mutant STING resulted in robust IRF and NF-κB activation, in addition to a marked production of cytokines, such as type I IFNs and IL-6, compared to WT STING transfection (Figure 5A and B). Notably, even at 10 µg/ml, HHMX strongly suppressed over-functional STING-mediated promoter activation and cytokine production, suggesting that HHMX may have a therapeutic potential for STING-related autoinflammatory diseases such as SAVI (Figure 5A and B). Furthermore, analysis of PBMCs from a SAVI patient carrying the STING N154S mutation (Figure S3A) by Western blotting revealed that PBMCs from SAVI patients exhibited elevated TBK1 and STAT1 phosphorylation in the absence of stimulation compared to that in healthy control PBMCs. Indeed, HHMX treatment suppressed the spontaneous TBK1 and STAT1 phosphorylation observed in SAVI patient PBMCs, suggesting that HHMX may have therapeutic potential for SAVI. (Figure 5C and S3A).

**Figure 5:**
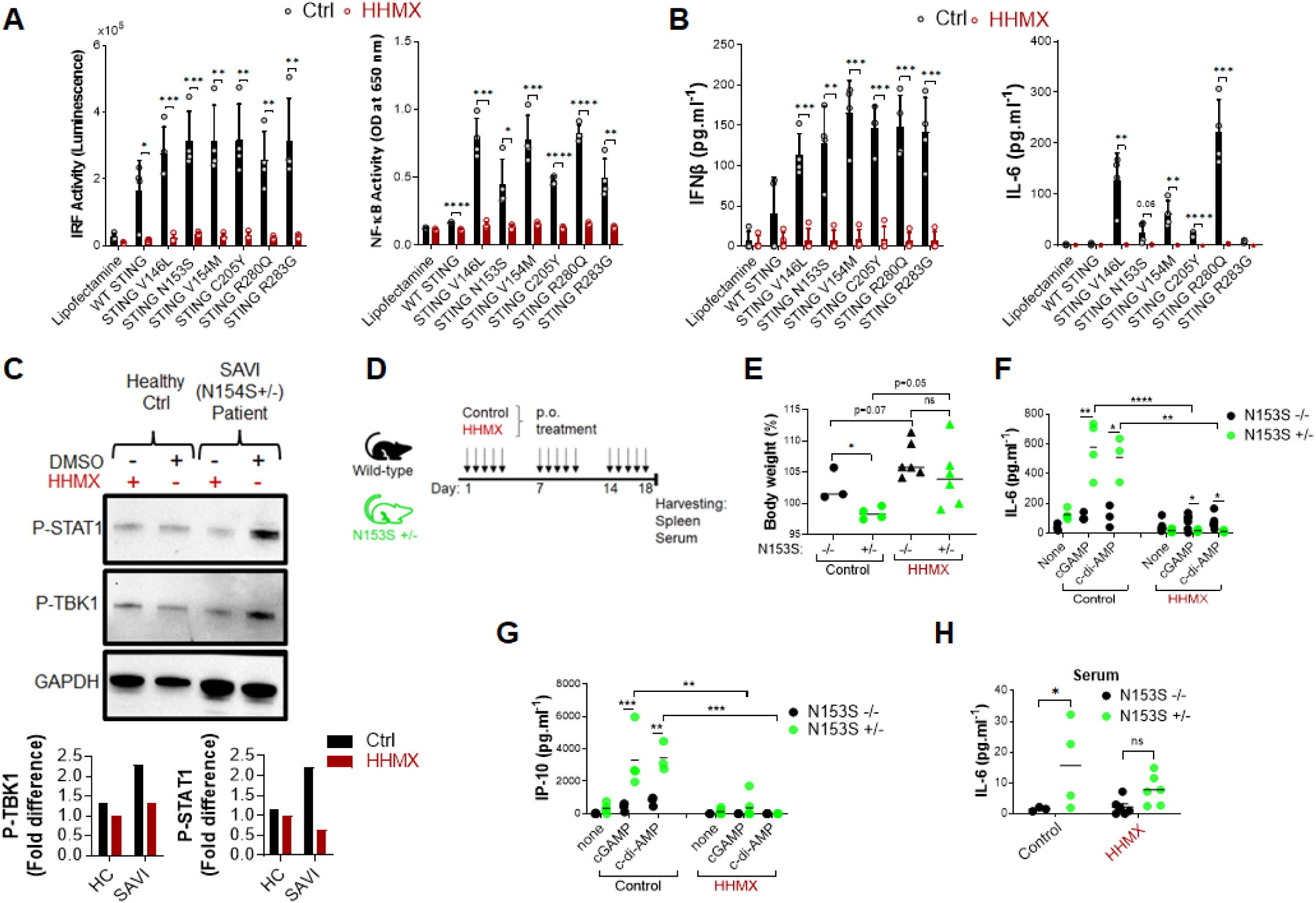
HHMX shows therapeutic potential against the STING-mediated autoinflammatory disease SAVI. **(A)** J774 dual reporter cells were transfected with the plasmids expressing WT STING or STING containing V146L, N153S, V154M, C205Y, R280Q, or R283G mutations together with HHMX (10 µg/ml) co-stimulation for 24 h in four independent experiments. IRF activity was determined by measuring secreted luciferase activity in the supernatants. NF-κB activity was determined by measuring secreted embryonic alkaline phosphatase (SEAP) activity in the supernatant. Data are shown as the mean ± SD of four independent experiments (n=4) (*p < 0.05, **p < 0.01, ***p < 0.001, ****p < 0.0001, Student’s t test.) **(B)** The concentrations of IFNβ and IL-6 in the supernatants were measured using ELISA. Data are shown as the mean ± SD of four independent experiments (n=4) (**p < 0.01, ***p < 0.001, ****p < 0.0001, Student’s t test.) **(C)** Fresh human PBMCs isolated from a healthy donor or SAVI patient bearing heterozygous STING N154S mutation were treated with HHMX (30 µg/ml) for 24 h and after protein isolation; the phosphorylation of TBK1 and STAT1 was investigated using western blotting. Graphs showing the band quantifications for P-TBK1 and P-STAT1 after normalization to GAPDH were plotted. **(D)** Eight–to-thirteen-week-old N153S^+/-^ SAVI mice (n=4-6) and their WT littermate (n=3-6) controls were orally administered 500 µg of HHMX or control, respectively, for 3 weeks every weekday (5 times/week). **(E)** Body weight and symptoms of mice were monitored until the day of sacrifice. Body weight percentage on day 18 are shown as dot plots with mean values (*p < 0.05, Student’s t test). **(F and G)** Splenocytes isolated from the control or HHMX treatment mice were stimulated with cGAMP and c-di-AMP for 24 h and the supernatant **(F)** IL-6 and **(G)** IP-10 levels were measured using ELISA. Data is the representative of two independent experiments and individual data from each mouse are shown (*p < 0.05, **p < 0.01, ***p < 0.001, ****p < 0.0001, Student’s t test). **(H)** The serum IL-6 levels of the HHMX-treated or control mice were measured using ELISA with the serum samples taken on the day of sacrifice. Data is the representative of two independent experiments and individual data from each mouse are shown (*p < 0.05, Student’s t test).

Based on the evidence regarding the potential therapeutic effect of HHMX in SAVI patient blood, we next tested the therapeutic potential of HHMX in SAVI mice bearing heterozygous STING N153S mutation (Figure S3B). Briefly, 8–13-week-old N153S^+/-^ SAVI mice generated by CRISPR/Cas9 technology, and their WT littermate controls were orally administered 500 µg of HHMX or vehicle for 3 weeks every weekday (5 times/week). The body weight and symptoms of the experimental mice were monitored until the day of sacrifice (Figure 5D). In the control treatment group, SAVI mice were found to lose weight after 2 weeks of treatment, with their weights being significantly lower than those of WT mice by day 18. Surprisingly, HHMX-treated SAVI mice did not show any weight loss or disease symptoms during the experiment period (Figure 5E). Furthermore, the stimulation of the splenocytes from the control treatment group mice with STING agonists cGAMP and c-di-AMP showed that SAVI mice had significantly elevated IL-6 and IP-10 responses in response to STING ligand stimulation compared to those from WT mice. By contrast, HHMX-treated SAVI mice showed significantly decreased IL-6 and IP-10 responses to STING agonist stimulation compared to those in WT mice (Figure 5F and G). Moreover, SAVI mice in the control treatment group showed significantly elevated serum IL-6 levels compared to those in the WT mice in the control treatment group (Figure 5 H). Moreover, there was no significant difference between the WT and SAVI mice in the HHMX treatment group, indicating that HHMX suppressed IL-6 elevation in SAVI mice (Figure 5H). Collectively, these results indicate that HHMX has a strong therapeutic potential for STING-dependent autoinflammatory diseases such as SAVI.

## Discussion

Accumulating evidence has indicated that DMXAA is a STING agonist that can exert potent adjuvant and anti-tumor activities in mice, but not in humans due to its inability to activate the human STING pathway (5,8,22,23). However, DMXAA has shown partial anti-tumor activity in humans as it has been studied in NSCLC patients up to Phase III (15). Although the multi-kinase inhibitory activity of DMXAA reported in human umbilical vein endothelial cells may partially explain this partial anti-tumor effect observed in humans, there are no studies on the suppressive effects of DMXAA on the human STING pathway, which may further explain this partial anti-tumor effect; chronic innate immune activation has been reported to induce tumorigenesis in a STING-dependent manner, with STING activation showing protumorigenic activity in lung cancer cells with a low antigenicity due to STING-mediated IDO activation in the tumor microenvironment (18,24). Herein, for the first time, we propose DMXAA as a suppressor of STING-induced immune responses in humans. At relatively higher doses, which was reported to be the optimal dose for type I IFN production from mouse peritoneal macrophages (25), DMXAA was found to suppress STING-induced IRF, NF-κB, type I IFN, and IL-6 responses by acting as a partial human STING agonist, interfering with agonistic STING activation. In addition, the novel DMXAA derivative HHMX more strongly antagonized STING pathway activation than DMXAA and became a promising therapeutic agent for STING-associated autoinflammatory diseases, being found to suppress over-functional STING (SAVI mutants)-induced immune responses *in vitro*, not only in cell lines but also in hPBMCs from SAVI patients, in addition to showing potent therapeutic effects in our SAVI mouse model by mitigating disease progression (Figure 6).

**Figure 6.**
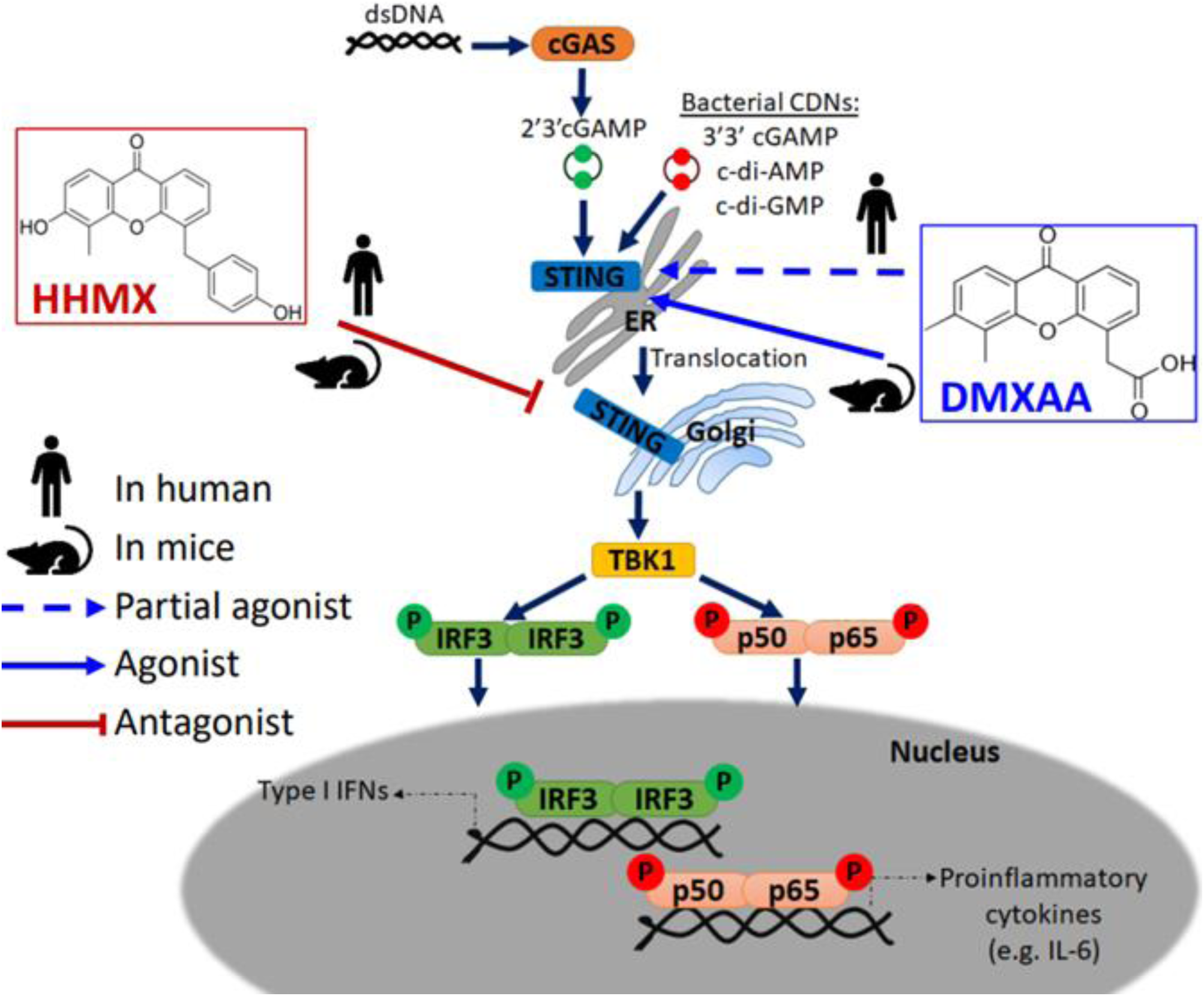
Mode of action of DMXAA and its derivative HHMX in human and mice. Mouse-specific STING agonist DMXAA is a partial STING agonist interfering with agonistic STING activation in humans. Based on this data, we developed a novel DMXAA derivative, 3-hydroxy-5-(4-hydroxybenzyl)-4-methyl-9H-xhanthen-9one (HHMX), which could antagonize STING signaling pathway both in human and mice to become potential therapeutic agent for STING-associated autoinflammatory diseases, like SAVI.

Our *in vitro* data using hPBMCs indicated that DMXAA exerts a greater suppressive activity on the STING-IRF3-type I IFN axis than on the STING-NF-κB-IL-6 axis (Figure 1). In agreement with these results, Zhong et al. also reported that STING could induce type I IFN production via IRF3, but not NF-κB, suggesting that pathways inducing type I IFNs and IL-6 could be regulated differentially (18). Importantly, this suppressive effect is not due to DMXAA-mediated cell death, as it may also influence cytokine production. Moreover, western blot data indicated that DMXAA showed a significant enhancing effect on STING-induced IRF3 and TBK1 phosphorylation rather than showing a suppressive effect, which suggests that DMXAA could be a partial human STING agonist interfering with agonistic STING pathway activation, since relatively higher doses of DMXAA (wherein a ratio of 1:10 agonist:DMXAA) are required to observe its suppressive effect and the partial agonists are defined as the compounds that can bind to their receptors but fail to fully activate them unlike conventional/true agonists (26). Rather partial agonists show antagonistic effect when they are used at higher doses due to competition of those partial agonists with the other conventional/true agonists, which is in alignment with our data about DMXAA and STING agonists, like CDNs. Nevertheless, it is also possible that post-transcriptional mechanisms acting downstream of TBK1, IRF3, or NF-κB phosphorylation events may be responsible for the suppressive effect of DMXAA on the human STING pathway. Furthermore, differential responses of THP1 dual reporter cells to different STING agonists were observed in our study with c-di-AMP and c-di-GMP being weaker activators of STING than cGAMP both for TBK1, IRF3 phosphorylation and stimulation of IRF and NF-κB activities, as well as cytokine production (Figure 1B-D, S2C and figure 3 C-D). Indeed, Tsuchiya et al. also reported that optimal activation of STING upon agonist binding requires confirmational re-organization of its lid structure from open to close form and subsequent stabilization of this closed lid structure by the C-terminal tail to induce TBK1-mediated robust type I IFN production, which could be achieved by cGAMP but not by c-di-GMP due to inability of c-di-GMP to sustain this closed lid structure for optimal STING phosphorylation and type I IFN production (27). By contrast, differential IL-6 responses to DMXAA pretreatment between hPBMCs and human macrophages, as shown in figures 1 and 3, could result from the polymorphic nature of STING (21), differences in the type of cells responsive to STING agonist in hPBMCs, or existence of humoral factors from specific types of cells in hPBMCs affecting other cells, which complicates the analysis of the suppressive effect of DMXAA in hPBMCs. Therefore, to analyze these mechanisms, we focused on human cell lines rather than PBMCs.

Our hPBMC experiments directly comparing the suppressive effects of DMXAA and HHMX showed that HHMX was a stronger antagonist of the STING pathway, most likely acting on molecules downstream of STING, such as TBK1, unlike DMXAA, because it could also suppress Poly I:C-mediated immune responses that were induced via STING-independent but TBK1-dependent mechanisms (Figure 3F and G, Figure 6). Furthermore, unlike DMXAA, HHMX antagonizes the STING pathway independent of STING polymorphism, as it showed a potent suppressive effect on all hPBMCs tested when used at the same dose as DMXAA, supporting our hypothesis that HHMX acts on molecules downstream of STING (Figure 1A and 3B). However, there still is a possibility that HHMX may directly bind to STING and antagonize its activity.

Using *in vitro* experiments, we demonstrated that HHMX can potently suppress exacerbated IRF, NF-κB, and cytokine responses induced by over-functional STING with various SAVI mutations, indicative of the therapeutic potential of HHMX as a drug for STING-mediated autoinflammatory diseases (Figure 5A and B). Importantly, HHMX was found to suppress the elevated STAT1 and TBK1 phosphorylation levels observed in PBMCs (Figure 5C). Indeed, because JAK-STAT inhibitors showed a similar suppressive effect on over-functional STING-induced immune responses, we hypothesize that STAT1 (downstream of STING-IRF3-type I IFN axis), which is constitutively phosphorylated in B and T cells of certain SAVI patients, may act as another potential target, but requires further investigation for the antagonistic binding of HHMX, in addition to TBK1 (19). Nevertheless, HHMX prevented the development of SAVI in the SAVI mouse model bearing the heterozygous STING N153S mutation, suggesting that HHMX may function as a potent immunotherapeutic agent to prevent the exacerbation of disease progression (Figure 5E–H). On the other hand, therapeutic effect of DMXAA derivatives, such as HHMX, is yet to be tested in other diseases that are thought to be dependent, at least in part, on STING. Moreover, one of the challenges in developing STING-based therapeutic agents for STING-associated autoinflammatory diseases is the variability of responsiveness to STING ligands among individuals due to the highly polymorphic nature of STING (21). However, because our data indicated that HHMX antagonizes downstream molecules of STING such as TBK1 or STAT1, rather than STING, polymorphism is not an issue for its therapeutic application for autoinflammatory diseases.

In conclusion, our study revealed the unanticipated suppressive effect of DMXAA on the human STING pathway, most likely due to interference of higher doses of DMXAA with agonist binding to STING. In addition, we introduced a novel DMXAA derivative HHMX, which acts as a potent antagonist for STING pathway activation via mechanisms involving the action of HHMX on downstream molecules of STING such as TBK1 (Figure 6). Taken together, our data provide further insights into the mechanisms of action of the mouse-selective STING agonist DMXAA in human cells and suggest that DMXAA derivatives such as HHMX are potential therapeutic agents for STING-associated autoinflammatory diseases, including SAVI.

## Materials and Methods

### Cell lines and reagents

THP1 dual reporter cells (catalog # thpd-nfis), HEK-Blue IFNα/β reporter cells (catalog # hkb-ifnabv2) and J774 dual reporter cells (catalog # j774d-nfis), were purchased from InvivoGen (San Diego, CA, USA). THP1 dual reporter cells were maintained in RPMI (Nacalai Tesque Inc.,Kyoto, Japan) supplemented with 10% FCS (Sigma-Aldrich, Missouri, USA), 1% penicillin/streptomycin (Nacalai Tesque Inc.,Kyoto, Japan), blasticidin (InvivoGen, catalog # ant-bl-05) (10 µg/ml), and zeocin (InvivoGen, catalog # ant-zn-05) (100 µg/ml). J774 dual reporter cells were maintained in DMEM (Nacalai Tesque Inc.,Kyoto, Japan) supplemented with 10% FCS, 1% penicillin/streptomycin, blasticidin (5 µg/ml), and zeocin (100 µg/ml). HEK-Blue IFNα/β reporter cells were maintained in DMEM supplemented with 10% FCS, 1% penicillin/streptomycin, blasticidin (30 µg/ml), and zeocin (100 µg/ml). All transfections were performed using Lipofectamine 2000 (Invitrogen, Massachusetts, USA).

DMXAA derivatives were synthesized and provided by the UBE Corporation (Japan) and they were dissolved in dimethylsulfoxide (DMSO). 2′3′ cGAMP (catalog # tlrl-nacga23-02), 3′3′cGAMP (catalog # tlrl-nacga), R848 (catalog # tlrl-r848), PolydA:dT (catalog # tlrl-patn), and Poly I:C (catalog # tlrl-pic) were purchased from Invivogen and they were dissolved in endotoxin/nuclease-free water. c-di-AMP was kindly donated by Yamasa (Chiba, Japan). K3 CpG ODN was synthesized at GeneDesign, as previously described (28). Both K3 CpG and c-di-AMP were dissolved in endotoxin/nuclease-free water. DMXAA was purchased from Sigma-Aldrich (St. Louis, MO, USA, catalog # D5817) and was dissolved in DMSO. P-TBK1/NAK (Ser172, catalog # 5483), TBK-1/NAK (catalog # 3013), P-NF-Κb p65 (Ser536, catalog # 3033), NF-κB p65 (catalog # 8242), P-IRF3 (Ser396, catalog # 29047), IRF3 (catalog # 4302), and β-tubulin (catalog #86298) antibodies were purchased from Cell Signaling Technology (Danvers, MA, USA).

Constructs expressing WT STING (pcIneo mSTING-HA) were kindly provided by Dr. Nao Jonai (National Institutes of Biomedical Innovation, Health and Nutrition). Constructs expressing V146L, N153S, V154M, C205Y, R280Q, or R283G mutant forms of STING were prepared using site-directed mutagenesis and PCR amplification via the primers below and DpnI digestion:

V146L Forward: 5′-GAAGTCTCTGCACTCTGTGAAGAAAAGAAG-3′
V146L Reverse: 5′-CTTCTTTTCTTCACAGAGTGCAGAGACTTC-3′
N153S Forward: 5′-AGAAAAGAAGTTAAGTGTTGCCCACGGGCT-3′
N153S Reverse: 5′-AGCCCGTGGGCAACACTTAACTTCTTTTCT-3′
V154M Forward: 5′-AAAGAAGTTAAATATGGCCCACGGGCTGGC-3′
V154M Reverse: 5′-GCCAGCCCGTGGGCCATATTTAACTTCTTT-3′
C205Y Forward: 5′-TCTTTCCATTGGACTATGGGGTGCCTGACA-3′
C205Y Reverse: 5′-TGTCAGGCACCCCATAGTCCAATGGAAAGA-3′
R280Q Forward: 5′-AGCTGGCTTCAGTCAGGAGGATCGGCTTGA-3′
R280Q Reverse: 5′-TCAAGCCGATCCTCCTGACTGAAGCCAGCT-3′
R283G Forward: 5′-CAGTCGGGAGGATGGGCTTGAGCAGGCTAA-3′
R283G Reverse: 5′-TTAGCCTGCTCAAGCCCATCCTCCCGACTG-3′

### Measurement of NF-κB and IRF activity in THP1 or J774 dual reporter cell lines

THP1 dual reporter cells were differentiated using 100 ng/ml PMA for 3 h and then washed with PBS. Three days after culturing in RPMI, the cells were washed with PBS again and stimulated/transfected in RPMI for the indicated time points in the figure legends. J774 cells were stimulated/transfected in DMEM for the indicated time points in the figure legends.

After stimulation at a density of either 5 × 10^5^ THP1 dual reporter cells/ml (in 200 µl) or 2.8 × 10^5^ J774 dual reporter cells/ml (in 200 µl) in 96-well plates, the supernatants of the cells were incubated with QUANTI-Blue (InvivoGen, catalog # rep-qbs) for 1–6 h, and the optical density (OD) at 650 nm was measured to determine SEAP activity, representative of NF-κB activity. To determine IRF activity, 20 µL of the supernatant was mixed with 50 µL of QUANTI-Luc (InvivoGen, catalog # rep-qlc1), and luminescence was measured using a microplate reader (Powerscan HT).

### Real Time-PCR analysis

RNA isolation from the cultured THP1 dual reporter cells was performed by using ReliaPrep RNA Cell Miniprep Kit (Promega, USA, catalog # Z6011) according to the instructions of the manufacturer. After the isolation, Nanodrop Spectrophotometer (ND-1000) was used to assess RNA quality. Genomic DNA removal was achieved by incubating 0.5 µg of the isolated RNA with DN Master Mix provided by the ReverTra Ace qPCR RT Master Mix with gDNA Remover kit (TOYOBO, Japan, catalog # FSQ-301) for 5 min at 37 °C. After genomic DNA removal, reverse transcription reaction was performed by adding RT Master Mix II (provided by the same kit) into the reaction mix and incubating it at 37 °C for 15 min and 98 °C for 5 min in the Veriti Thermal Cycler (Applied Biosystems, USA). Quantitative analysis of *IL-6* and *IFNB* expression in THP1 dual reporter cells was performed by StepOne Real-Time PCR Systems (Thermo Fisher Scientific) by using 18S rRNA for normalization. 2^-ΔΔCt^ formula was used for calculation of relative changes in the expression of *IFNB* and *IL-6* for each stimulation based on control treatment. Particularly, following formula was used to calculate relative expression values for *IFNB* and *IL-6*: 2^-((Ct _treatment_ of IL-6/IFNB)-(Ct _treatment_ of 18S)-(average(Ct _control treatment_ of IL-6/IFNB)-(Ct _control treatment_ of 18S))).

*<u>Primers used for qPCR:</u>*

rRNA Forward: AAACGGCTACCACATCCAAG
rRNA Reverse: CCTCCAATGGATCCTCGTTA
IFN-b Forward: AGCACTGGCTGGAATGAGAC
IFN-b Reverse: CTATGGTCCAGGCACAGTGA
IL-6 Forward: AGTTCCTGCAGAAAAAGGCA
IL-6 Reverse: AAAGCTGCGCAGAATGAGAT

### Human PBMC experiments

All hPBMC experiments were performed with the approval of the Institutional Review Board of the Institute of Medical Science, the University of Tokyo (IMSUT) (approval number: 2019-25-0919). All procedures involving human samples adhered to the World Medical Association (WMA) Declaration of Helsinki and The Department of Health and Human Services Belmont Report.

Department of Health and Human Services Belmont Report Frozen PBMCs, purchased from Lonza (Basel, Switzerland), were prepared and kindly provided by Takato Kusakabe. hPBMCs from healthy blood donors were isolated using Ficoll-Paque Plus (GE Healthcare, Little Chalfont, UK). PBMCs (1 × 10^6^) were cultured in RPMI containing 10% FCS and 1% penicillin/streptomycin. After 90 min of DMXAA or HHMX pretreatment, the hPBMCs were stimulated with CDNs (10 µg/ml) or PolydA:dT (1 µg/ml) for 24 h. IFNγ and IL-6 production was measured using ELISA. Type I IFN production was measured using HEK-Blue IFNα/β reporter cells or ELISA.

The SAVI patient bearing the STING N154S mutation was a pediatric patient in Turkey, and the required samples were obtained by Prof. Seza Ozen from Hacettepe University in Turkey after obtaining informed consent from the parents of the patient. This SAVI patient was treated with a Jak1/2 inhibitor (Baricitinib). The study protocol was approved by the Institutional Review Board of Hacettepe University (IRB no. 16969557-2258, project no. GO 19/1003 and IRB approval no at IMSUT: 2022-5-0519).

### Animal studies

Animal protocols were approved by the Animal Resource Center for Infectious Diseases, RIMD and IFReC, Osaka University (approval no. 14012); the Faculty of Medicine, Osaka University (approval no. 280001); and the animal facility of the Institute of Medical Science, University of Tokyo (IMSUT) (approval no. PA19-63).

C57BL/6 mice were maintained in pathogen-free animal facilities at the Animal Resource Center for Infectious Diseases, RIMD and IFReC, Osaka University, and the Faculty of Medicine, Osaka University. Mice were fed a standard diet and water. STING N153S^+/-^ KI mice were generated by Prof. Yamamoto using CRISPR genome editing technology as previously described (29, 30). WT male or female mice were crossed with heterozygous mice to generate heterozygous mice. WT littermates co-housed with heterozygous mice were used as the WT control mice for this study. HHMX was dissolved in DMSO and diluted with PBS for the p.o. treatment. Eight–to-thirteen-week-old N153S^+/-^ SAVI mice and their WT littermate controls were orally administered 500 µg of HHMX and DMSO control, respectively, for 3 weeks every weekday (5 times/week). Body weight and symptoms were monitored throughout the study until sacrifice. Upon sacrifice, the splenocytes were stimulated with 10 µg/ml of cGAMP or c-di-AMP for 24 h, and the supernatant cytokine levels were measured using ELISA.

### Genotyping

Ear tissues obtained from mice were digested with proteinase K, and DNA was isolated and purified using the Wizard SV Genomic DNA Purification System (Promega, WI, USA, catalog # A2920) according to the manufacturer’s protocol. Two separate PCRs were performed for genotyping, as follows:

<u>Primers for WT allele detection:</u>

Forward primer: 5′-GCCTCGCACGAACTTGGACTACTGT-3′
WT reverse primer: 5′-CAGCCCGTGGGCAACATT-3′

<u>Primers for STING N153S mutant allele detection:</u>

Forward primer: GCCTCGCACGAACTTGGACTACTGT
STING N153S reverse primer: CAGCCCGTGGGCAACAGA

Primers recognizing sequences in exon 5 of Tmem173 (STING) were added to the PCR buffer for KOD-Plus-Neo (TOYOBO) at a final concentration of 0.3 μM. PCR was performed with an initial denaturation step at 94°C for 2 min, followed by 30 cycles of denaturation at 98°C for 10 s, and annealing and extension at 68°C for 25 s. Mice positive for the STING N153S mutation were identified by the appearance of a 700-bp band using agarose gel electrophoresis and sequencing (Figure S3B).

### In vitro cytotoxicity assay and DNA release measurements

Supernatants from stimulated cells were mixed with the substrate mix at a 1:1 ratio and incubated for 15 min at room temperature. The percentage cytotoxicity was calculated using the OD measurements at 490 nm, according to the instructions of Non-Radioactive Cytotoxicity Assay Kit (Promega, catalog # G1780). The positive control (100% cytotoxicity) for the assay was prepared by incubating the cells with Triton X-100 at 37°C for 15 min.

DNA in the HHMX-stimulated supernatants of THP1 dual reporter cells was quantified using the QuantiFluor dsDNA Detection Kit from Promega (catalog # E2670) according to the manufacturer’s instructions.

### Cytokine measurement

Human IL-6 (catalog # DY206), human IFN-γ (catalog # DY285B), human IP-10 (catalog #DY266), mouse IL-6 (catalog # DY406), mouse IFNβ (catalog # 42400-2) cytokine levels were measured using ELISA kits from R&D Systems (Minneapolis, MN, USA). Human IFNβ levels were measured using an ELISA kit (PBL Assay Science, NJ, USA, catalog # 41410).

To determine the levels of type I IFNs in the stimulated human PBMCs, HEK-Blue IFNα/β reporter cells were stimulated with 20 µL of culture supernatants for 24 h. The resulting supernatants were incubated with QUANTI-Blue for 1–6 h and OD at 650 nm was measured to determine SEAP activity, representative of type I IFN levels in the culture supernatants.

Customized cytokine 25-Plex Human ProcartaPlex Assay (customized from Cytokine 25-Plex Human ProcartaPlex Panel 1B with the catalog # EPX250-12166-901, Thermo Fisher Scientific, USA) was used to measure multiple cytokines in the supernatants of stimulated human PBMCs.

### SDS-PAGE and western blotting

After the stimulation/transfection (transfection with lipofectamine 2000 (Thermo Fisher Scientific, catalog # 11668019)) of PMA-differentiated THP1 dual reporter cells, or J774 dual reporter cells in 24-well plates, the cells were washed with ice-cold PBS and directly lysed in 150 µl of sample loading buffer containing 0.1 M Tris/HCl (pH 6.7), 2% SDS, 10% glycerol, bromophenol blue, and 1% β-mercaptoethanol. Lysates were boiled at 95°C for 10 min and vortexed well before loading onto 10% SDS-PAGE gels. Proteins were transferred to PVDF membranes using an iBlot2 gel transfer device (Thermo Fisher Scientific, Massachusetts, USA). Next, the membranes were blocked in TBST containing 5% skim milk at room temperature for 1 h and incubated with the appropriate primary antibodies diluted in TBST containing 5% BSA at room temperature for 1 h. The membranes were then washed three times with TBST and incubated with HRP-linked anti-rabbit IgG antibody (Cell Signaling Technology, catalog # 7074) in TBST containing 5% BSA for 1 h at room temperature. Protein bands were visualized by treating the membrane with Immobilon Western Chemiluminescent HRP Substrate (Merck Millipore, Darmstadt, Germany, catalog # 42029053) and developing on the films using a KODAK X-OMAT 1000A Processor.

## Data Availability

This study includes no data deposited in external repositories.

## Statistical analysis

Statistical tests were performed if n ≥ 3with GraphPad Prism software (La Jolla, CA, USA). Student’s *t*-test was used for comparison of each group for in vivo experiments. Student’s *t*-test or one-way analysis of variance with the appropriate multiple comparison tests, such as Sidak’s test, were used for the statistical analyses for in vitro studies (*p < 0.05, **p < 0.01, ***p < 0.001, ****p < 0.0001) based on the distribution of the data.

## Author Contributions

B.T., T.S. and K.J.I. designed the study. B.T., and K.J.I. wrote the manuscript. B.T., T.S., K.H., N.S. and E.S. performed the experiments. M.Y. generated and provided the SAVI model mice for this study. All other authors have contributed to data collection and interpretation, and critically reviewed the manuscript.

## Acknowledgments

HHMX was synthesized and kindly provided by UBE Corporation (Japan). c-di-AMP and c-di-GMP were kindly provided by Yamasa (Chiba, Japan). We thank Mizuho Yamashita, Naomi Sato, Ayane Takao, Mai Onaga and Seigo Akaeda for their technical assistance.

## Data Availability Statement

All the data related to this study is presented in this manuscript. Additional data and raw data of the presented data can be provided by the authors upon request.

## References

1. Sun L, Wu J, Du F, Chen X, Chen ZJ. Cyclic GMP-AMP Synthase Is a cytosolic DNA sensor that activates tye type I IFN pathway. Science. 2013;339:786–91.

2. Mankan AK, Schmidt T, Chauhan D, Goldeck M, Höning K, Gaidt M, et al. Cytosolic RNA:DNA hybrids activate the cGAS–STING axis. EMBO J. 2014;33:2937–46.

3. McWhirter SM, Barbalat R, Monroe KM, Fontana MF, Hyodo M, Joncker NT, et al. A host type I interferon response is induced by cytosolic sensing of the bacterial second messenger cyclic-di-GMP. J Exp Med. 2009;206:1899–911.

4. Burdette DL, Monroe KM, Sotelo-Troha K, Iwig JS, Eckert B, Hyodo M, et al. STING is a direct innate immune sensor of cyclic di-GMP. Nature. 2011;478:515–8.

5. Gao P, Zillinger T, Wang W, Ascano M, Dai P, Hartmann G, et al. Binding-pocket and lid-region substitutions render human STING sensitive to the species-specific drug DMXAA. Cell Rep. 2014;8:1668–76.

6. Li XD, Wu J, Gao D, Wang H, Sun L, Chen ZJ. Pivotal roles of cGAS-cGAMP signaling in antiviral defense and immune adjuvant effects. Science. 2013;341(6152):1390–4.

7. Tang CK, Aoshi T, Jounai N, Ito J, Ohata K, Kobiyama K, et al. The Chemotherapeutic Agent DMXAA as a Unique IRF3-Dependent Type-2 Vaccine Adjuvant. PLoS One. 2013;8:6–11.

8. Woo S-R, Fuertes MB, Corrales L, Spranger S, Furdyna MJ, Leung MYK, et al. STING-Dependent Cytosolic DNA Sensing Mediates Innate Immune Recognition of Immunogenic Tumors. Immunity. 2014;41:830–42.

9. Deng L, Liang H, Xu M, Yang X, Burnette B, Arina A, et al. STING-dependent cytosolic DNA sensing promotes radiation-induced type I interferon-dependent antitumor immunity in immunogenic tumors. Immunity. 2014;41(5):843–52.

10. Corrales L, Glickman LH, McWhirter SM, Kanne DB, Sivick KE, Katibah GE, et al. Direct Activation of STING in the Tumor Microenvironment Leads to Potent and Systemic Tumor Regression and Immunity. Cell Rep. 2015;11:1018–30.

11. Barber GN. STING-dependent cytosolic DNA sensing pathways. Trends Immunol. 2014;35:88–93.

12. Liu Y, Jesus AA, Marrero B, Yang D, Ramsey SE, Sanchez GAM, et al. Activated STING in a Vascular and Pulmonary Syndrome. N Engl J Med. 2014;371:507–18.

13. Ahn J, Ruiz P, Barber GN. Intrinsic Self-DNA Triggers Inflammatory Disease Dependent on STING. J Immunol. 2014;193(9):4634–42.

14. Zhao L, Ching L-M, Kestell P, Baguley BC. The antitumour activity of 5,6-dimethylxanthenone-4-acetic acid (DMXAA) in TNF receptor-1 knockout mice. Br J Cancer. 2002;87(4):465–70.

15. Lara PN, Douillard JY, Nakagawa K, Von Pawel J, McKeage MJ, Albert I, et al. Randomized phase III placebo-controlled trial of carboplatin and paclitaxel with or without the vascular disrupting agent vadimezan (ASA404) in advanced non-small-cell lung cancer. J Clin Oncol. 2011;29(22):2965–71.

16. Kim S, Li L, Maliga Z, Yin Q, Wu H, Mitchison TJ. Anticancer Flavonoids Are Mouse-Selective STING Agonists. ACS Chem Biol. 2013;8:1396–401.

17. Ahn J, Xia T, Konno H, Konno K, Ruiz P, Barber GN. Inflammation-driven carcinogenesis is mediated through STING. Nat Commun. 2014;5:5166–75.

18. Zhong B, Yang Y, Li S, Wang YY, Li Y, Diao F, et al. The Adaptor Protein MITA Links Virus-Sensing Receptors to IRF3 Transcription Factor Activation. Immunity. 2008;29(4):538–50.

19. Melki I, Rose Y, Uggenti C, Van Eyck L, Frémond ML, Kitabayashi N, et al. Disease-associated mutations identify a novel region in human STING necessary for the control of type I interferon signaling. J Allergy Clin Immunol. 2017;140:543–52.

20. Gonda TA, Tu S, Wang TC, Gonda TA, Tu S, Chronic TCW, et al. Chronic inflammation, the tumor microenvironment and carcinogenesis. Cell cycle. 2009;8:2005– 13.

21. Yi G, Brendel VP, Shu C, Li P, Palanathan S, Cheng Kao C. Single Nucleotide Polymorphisms of Human STING Can Affect Innate Immune Response to Cyclic Dinucleotides. PLoS One. 2013;8:1–16.

22. Saito T, Gale M. Differential recognition of double-stranded RNA by RIG-I–like receptors in antiviral immunity. J Exp Med. 2008;205:1523–17.

23. Xiao-Dong Li, Jiaxi Wu, Daxing Gao, Hua Wang, Lijun Sun ZJC. Pivotal Roles of cGAS-cGAMP Signaling in Antiviral Defense and Immune Adjuvant Effects. Science. 2013;341:1390–5.

24. Buchanan CM, Shih J-H, Astin JW, Rewcastle GW, Flanagan JU, Crosier PS, et al. DMXAA (Vadimezan, ASA404) is a multi-kinase inhibitor targeting VEGFR2 in particular. Clin Sci. 2012;122(10):449–65.

25. Roberts ZJ, Goutagny N, Perera PY, Kato H, Kumar H, Kawai T, et al. The chemotherapeutic agent DMXAA potently and specifically activates the TBK1-IRF-3 signaling axis. J Exp Med. 2007;204(7):1559–69.

26. Zhu BT. Mechanistic explanation for the unique pharmacologic properties of receptor partial agonists. Biomed Pharmacother. 2005;59(3):76–89.

27. Tsuchiya Y, Jounai N, Takeshita F, Ishii KJ, Mizuguchi K. Ligand-induced Ordering of the C-terminal Tail Primes STING for Phosphorylation by TBK1. EBioMedicine. 2016;9:87–96.

28. Kobiyama K, Aoshi T, Narita H, Kuroda E, Hayashi M, Tetsutani K, et al. Nonagonistic Dectin-1 ligand transforms CpG into a multitask nanoparticulate TLR9 agonist. PNAS. 2014;111:3086–91.

29. Warner JD, Irizarry-Caro RA, Bennion BG, Ai TL, Smith AM, Miner CA, et al. STI NG-associated vasculopathy develops independently of IRF3 in mice. J Exp Med. 2017;214(11):3279–92.

30. Sasai M, Sakaguchi N, Ma JS, Nakamura S, Kawabata T, Bando H, et al. Essential role for GABARAP autophagy proteins in interferon-inducible GTPase-mediated host defense. 2017;18(8).

