## Supplementary Figures for "5,6-dimethylxanthenone-4-acetic acid (DMXAA), a Partial STING Agonist, Competes for Human STING Activation"

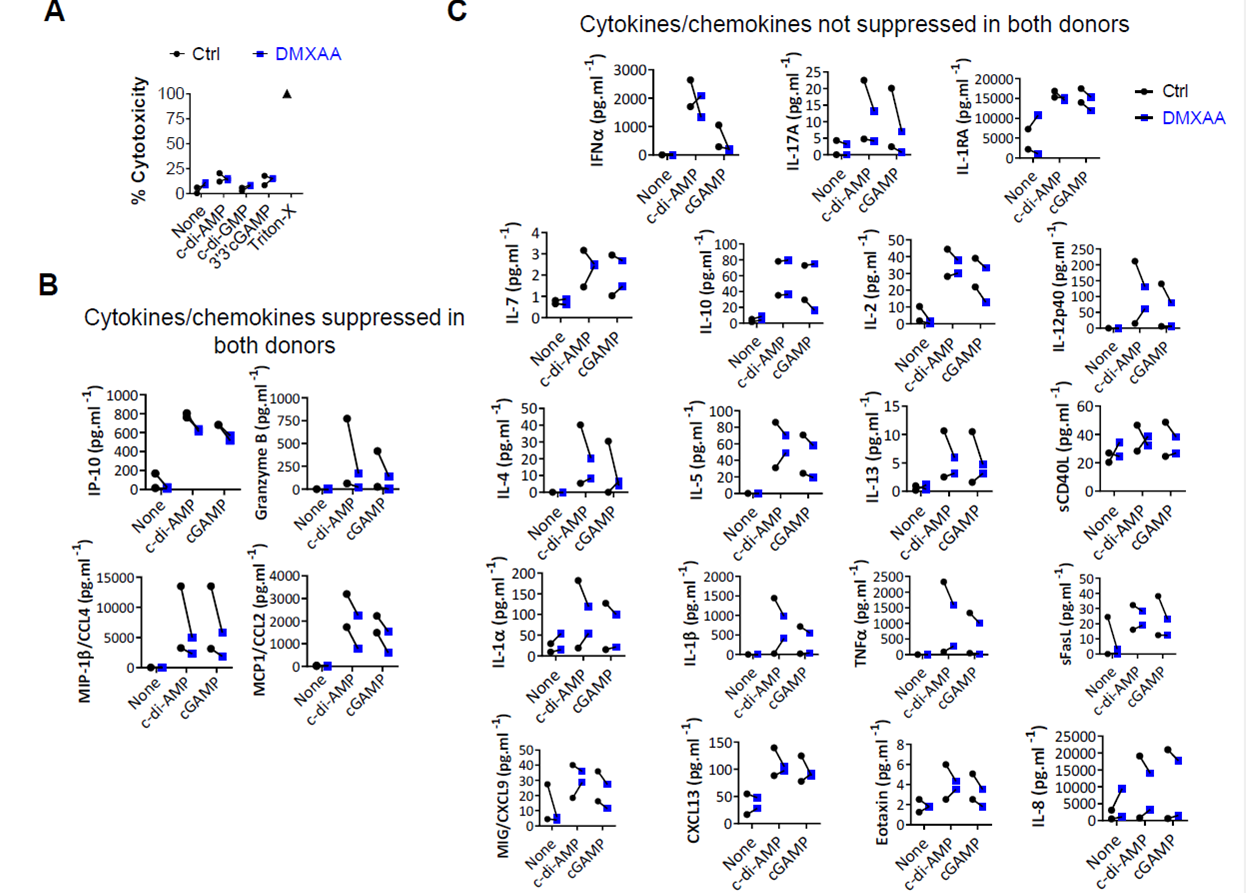

**Supplementary Figure 1.** Fresh human PBMCs from two healthy donors were stimulated with the indicated CDNs (10 µg/ml) for 24 h after 90 minutes of DMXAA (100 µg/ml) pretreatment. **(A)** Cell death was measured using lactate dehydrogenase (LDH) release assay in two of the PBMC donors. Data are shown as scatter plots showing individual values from each donor. **(B.C)** Cytokine production was measured using the human 25-plex Bio-plex Cytokine Assay. **(B)** IP-10, granzyme B, MIP-1b/CCL4, MCP1/CCL2 and **(C)** eotaxin, CXCL13, IL-1a, IL-1b, IL1-RA, IL-2, IL-4, IL-5, IL-6, IL-7, IL-8, IL-9, IL-10, IL-12p40, IL-13, IL-15, IL-17A, MIP-1a, MIG, MCP-1, TNFa, sCD40L and sFasL levels from Bio-plex analysis are shown as scatter plots showing individual values from each donor.

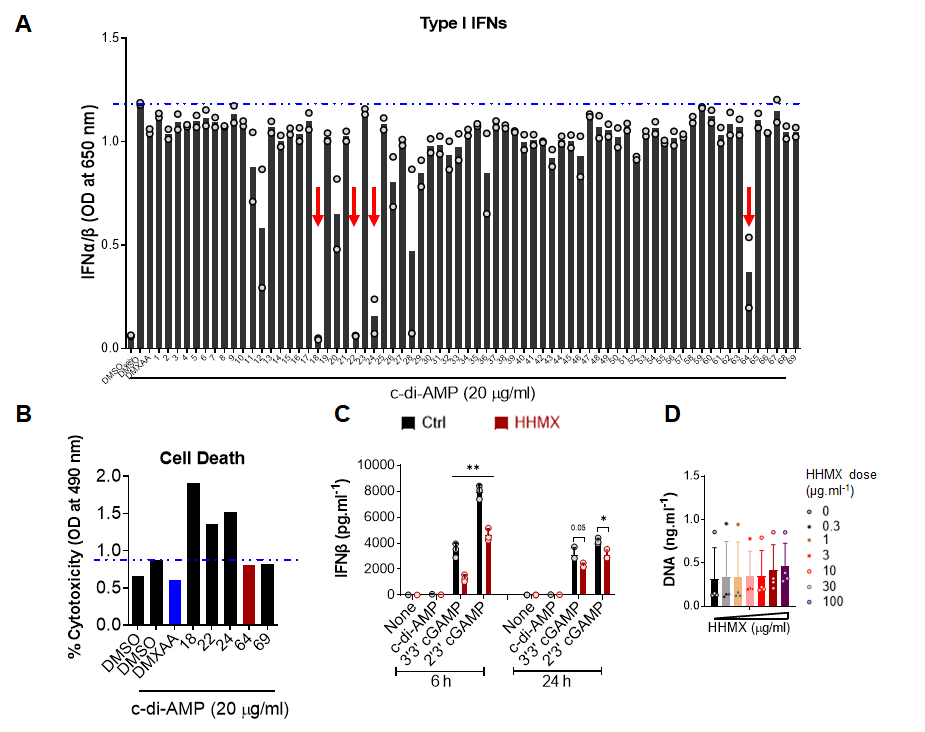

**Supplementary Figure 2: (A)** Fresh human PBMCs from two healthy donors were stimulated with c-di-AMP (20 µg/ml) with or without DMXAA or its derivatives (100 µg/ml) for 24 h. The levels of type I IFN in the supernatants were measured using HEK-Blue IFNα/β reporter cells. Bar graphs showing individual data and mean are plotted. Red arrows represent DMXAA derivatives with robust suppressive effect on STING-induced type I IFN production in human PBMCs. **(B)** Cell death was measured using the lactate dehydrogenase (LDH) release assay. Representative data from one PBMC donor among two different donors are shown as bar graph. **(C)** PMA-differentiated THP1 dual reporter cells were stimulated with HHMX (30 µg/ml) together with the indicated CDNs (10 µg/ml) for 6 or 24 h. The levels of type I IFN levels in the supernatants were measured using ELISA. Data are shown as the mean ± SD from three independent experiments (n=3) (*p < 0.05, **p < 0.01, Student’s t test). **(D)** PMA-differentiated THP1 dual reporter cells were stimulated with the indicated concentrations of HHMX (0, 0.3, 1, 3, 10, 30, 100 µg/ml) 24 h. DNA release was measured using the QuantiFluor dsDNA detection kit (Promega) according to manufacturer’s instructions. Data are shown as the mean ± SD from four independent experiments (n=4).

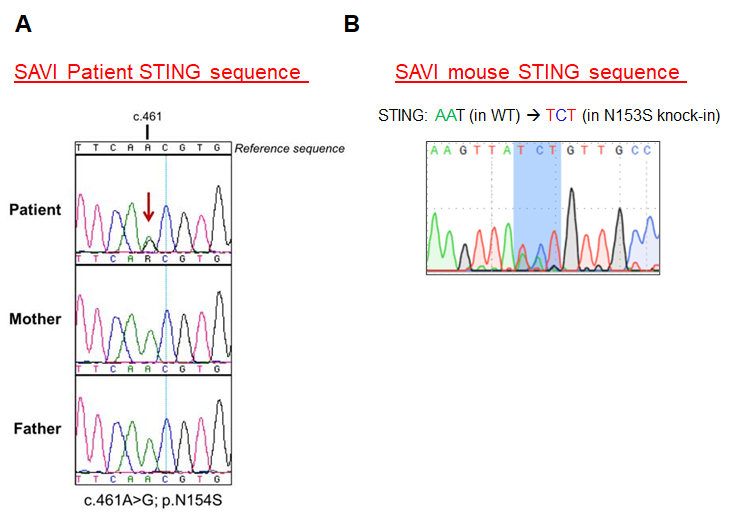

**Supplementary Figure 3: (A)** Sequencing data of STING in SAVI patients. **(B)** Sequencing data of STING in SAVI mouse.
